# Simulating the cellular context in synthetic datasets for cryo-electron tomography

**DOI:** 10.1101/2023.05.26.542411

**Authors:** Antonio Martinez-Sanchez, Lorenz Lamm, Marion Jasnin, Harold Phelippeau

**Affiliations:** Department of Information and Communications Engineering, Faculty of Computers Sciences, University of Murcia, Campus de Espinardo 30100 Murcia, Spain; Helmholtz Pioneer Campus, Helmholtz Munich, 85764 Neuherberg, Germany; Department of Chemistry, Technical University of Munich, 85748, Garching, Germany; Thermo Fisher Scientific, Bordeaux, France; Helmholtz AI, Helmholtz Munich, 85764 Neuherberg, Germany

**Keywords:** Cryo-Electron Tomography, Deep Learning, Image processing, Scientific Computation, Synthetic data generation

## Abstract

Cryo-electron tomography (cryo-ET) allows to visualize the cellular context at macromolecular level. To date, the impossibility of obtaining a reliable ground truth is limiting the application of deep learning-based image processing algorithms in this field. As a consequence, there is a growing demand of realistic synthetic datasets for training deep learning algorithms. In addition, besides assisting the acquisition and interpretation of experimental data, synthetic tomograms are used as reference models for cellular organization analysis from cellular tomograms. Current simulators in cryo-ET focus on reproducing distortions from image acquisition and tomogram reconstruction, however, they can not generate many of the low order features present in cellular tomograms.

Here we propose several geometric and organization models to simulate low order cellular structures imaged by cryo-ET. Specifically, clusters of any known cytosolic or membrane bound macromolecules, membranes with different geometries as well as different filamentous structures such as microtubules or actin-like networks. Moreover, we use parametrizable stochastic models to generate a high diversity of geometries and organizations to simulate representative and generalized datasets, including very crowded environments like those observed in native cells.

These models have been implemented in a multiplatform open-source Python package, including scripts to generate cryo-tomograms with adjustable sizes and resolutions. In addition, these scripts provide also distortion-free density maps besides the ground truth in different file formats for efficient access and advanced visualization. We show that such a realistic synthetic dataset can be readily used to train generalizable deep learning algorithms.

## I. Introduction

Cryo-electron tomography (cryo-ET) has emerged as a unique technique to generate tridimensional (3D) representations of the cellular context at sub-nanometric resolution [1]. However, volumetric images obtained by cryo-ET, called tomograms, suffer from high levels of noise and anisotropic distortions, mainly due to the missing wedge [2], which greatly complicate their analysis.

Due to its success in image processing and computer vision in recent years, deep learning (DL) algorithms, and especially convolutional neuronal networks (CNNs) [3], are being adapted to process cryo-ET tomograms at different steps of the workflow: reconstruction [4], denoising [5], [6], segmentation [7], [8] or macromolecular localization [9]. CNNs enable efficient analysis of local features in images because they take advantage of the inherent regular grid structure of digital images. When the training set is representative and large enough, CNN-based methods outperform classical approaches like template matching (TM) [10]. Nevertheless, the major bottleneck to apply DL in cryo-ET is the lack of ground truth. In most cases it is not possible to generate a reliable labeled dataset for a proper supervised training.

Regardless of the method used for segmenting cellular compartments and localizing macromolecules in cryo-tomograms, the next step is usually a quantitative analysis. In cryo-ET, statistical approaches are used to carry out macromolecular censuses [11], [12], analyze the distance between two proteins [13], study protein-membrane distance distribution [14] or prove the co-localization between two macromolecules [15]. The second-order statistics are more sophisticated [16], as they allow to analyze macromolecular organization on different scales, as well as merge and compare information coming from different tomograms. Second-order statistics have recently been used in cryo-ET to evaluate the clustering organization of some protein complexes under different functional states [17], show nanodomains defined by the co-localization of proteins bounded to two aligned membranes [18], describe the liquid-like organization of densely packed proteins by adjusting a statistical mechanics model [19]. In addition, rotations of macromolecules are typically analyzed to determine the rigidity of a bounding, e.g. between two macromolecules [20] or proteins and membranes [21]. Additionally, ad-hoc analysis can be defined to study filament networks [22] and membranes organization [23], [24]. In general, these statistical analyses require a synthetic null-model working as reference [16], which also allows to provide statistical confidence.

Cryo-ET data acquisition is costly as it requires high-end equipment and a lot of human expertise. Unfortunately, determining the applicability of a dataset for studying a specific structure is only possible after tomogram reconstruction, the last step in the workflow. Synthetic data can be used to streamline imaging parameters as well as to ensure sample appropriateness for solving a specific biological question. Simulations have already been used for particle picking and subtomogram averaging assessment [25], [26]. Recently, a tool specifically designed for membrane-embedded proteins has been proposed [27], however, it is limited to a single spherical membrane with uniformly distributed isolated proteins. Simulated annealing and molecular dynamics are used afterward to improve the packing and avoid volume overlapping by approximating molecules to spheres [26].

As shown above, simulating synthetic data containing lower-order structures present in the cellular environment will contribute greatly to the successful application of DL, to analyze quantitatively the organization of structures in cryo-ET, and assist the acquisition and interpretation of experimental data. There are already computational solutions for generating artificial data in transmission electron microscopy (TEM), but they focus on modelling the physics of image formation from atomic representations of macromolecules [28]–[30]. Methods for simulating cellular environments represent either isolated structures or a mixture of purely random distribution of macromolecules with some spherical vesicles [26], [31], [32]. In [33] some membrane-bound proteins are modelled but just by simple blobs in 2D. A concurrent pre-print [34] generates a more diverse scenario than previous approaches, which includes vesicles and membrane-proteins but not curved filaments. More importantly, this pre-print shows that coarse synthetic data already allows to train usable CNN models.

Here, we focus on producing richer contextual information, and directly generate synthetic 3D density maps, also known as phantoms, containing membranes with different combinations of local curvatures, clusters of macromolecules (either cytosolic or membrane-bound), as well as polymers such as microtubules or actin-like networks. Together with these clean 3D images, motif lists in different formats are provided as ground truth, allowing the generation of micrographs using any external TEM simulator to simulate the distortions introduced by the microscope and 3D reconstruction methods. Moreover, the introduction of generative neural networks holds the promise to produce datasets with more realistic noise and distortion [35], they can reproduce noise and distortions directly from the synthetic micrographs [36], but this is outside the scope of this study. In addition, techniques such as data augmentation [37], knowledge transfer [38] or contrastive learning [32] may allow to train DL-based algorithms without having a perfect representation of the experimental data. For statistical organization analysis, null-models do not require simulating data noise and distortions.

The simulation of high-order cellular structures such as organelles (mitochondrion, centriole, …) or membranes associations (synapses, Golgi, …) is beyond the scope of this paper. Here we focus on low-order structural features present in almost all cellular contexts. Low-order features are independent of each other and can thus be generated from parametric mathematical models to control their geometry and organization. In addition, we use parametrizable stochastic models to generate a high diversity of different combinations of geometries and organization to simulate representative and generalized datasets.

The rest of this paper is organized as follows: First, the developed methods are described, we detail how to model the geometry of each structure and their organization, and we explain how these models were implemented. Then, we show examples of synthetic data generated for replicating the cellular context in cryo-ET and use these data to train a generalized deep learning algorithm for semantic segmentation. We also evaluate certain properties of the proposed algorithms. Finally, the proposed methods and the obtained results are discussed.

## II. Methods

In this section, we first explain the mathematical models used to parameterize the cellular structures. Second, we describe the algorithms used to generate the synthetic data.

### A. Structural Models

The cellular context is represented by a density map *D*(x) : *𝒱 ⊂* ℝ^3^ ⟼ ℝ being x *∈* ℝ^3^ a point and *𝒱* the volume of interest (VOI), typically a cuboid whose dimensions are defined by the output tomogram, but in practice it can have any arbitrary shape.

1. *Membranes:* Membranes are modelled as parametric surfaces with double Gaussian profile along their local normal. The purpose of this profile is to simulate the lipid bilayer, which, in the TEM, looks like two parallel and very close electron-dense sheets. Although all biological membranes share the same structure, small visual differences may appear due to their distinct molecular composition or dissimilar imaging conditions. To account for these variations, the membrane profile is controlled by two parameters: membrane thickness or distance between both layers, *t*, and the layer variance, *σ*_*l*_, which controls the thickness of the layers. Below, we describe the parametric surfaces used to model the membrane geometries and how the double-layered profile is incorporated into them. **Sphere:** the outer, *L*_*o*_, and inner, *L*_*i*_, layers are defined by

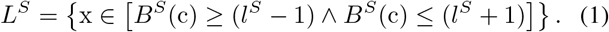

Where *B*^*S*^(c) = ∥x − c∥, c = (*c*_*x*_, *c*_*y*_, *c*_*z*_) is the sphere center, 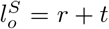 and 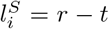 are the values of l for each layer, and *r* as the sphere radius. **Ellipsoid:** As above, the ellipsoidal layers are defined as

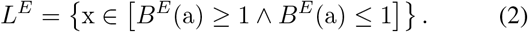

Where 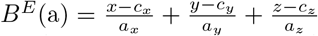, with a= (*a*_*x*,_ *a*_*y*,_ *a*_*z*_) the ellipsoid semi-axis lengths, for each layer the actual values for these lengths are a_*o*_ = a + *t* and a_*i*_ = a − *t*. **Torus:** Given a point *x* = (*x, y, z*) *∈* ℝ ^3^ the two toroid layers are defined by

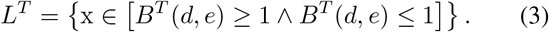

Where 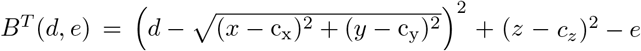, with *d* the major torus radius and *e* the minor torus radius, which have different values for each layer: *e*_*o*_ = *e* + *t* and *e*_*i*_ = *e* − *t*. The layers *L* described above have a non-smooth profile (hard edge). A 3D Gaussian filter 𝒢 is thus applied to obtain a smoothed version *L*^𝒢^ *∈ C*^2^ using the layer thickness *σ*_*l*_ defined above:

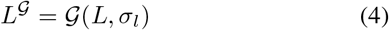

In addition to representing membranes as scalar fields, which is useful to generate the output density map, it is convenient to also represent them using 2-manifolds (surfaces) as

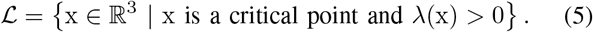

Being λ the largest eigenvalue of the Hessian Matrix of the scalar field 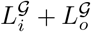 at point x. Since ℒ is the surface in the middle of both the inner and outer layers of the membrane, it can be used as the reference to localize membrane-bound macromolecules and to measure certain geometric properties of the membrane, such as areas or local curvatures.
2. *Filaments:* Filaments are modelled as flexible helical polymers. We consider a polymer as a chain of structural units. A structural unit is defined as a density function in 3D *M*(x) : ℝ^3^ → ℝ. The normalized tangent vector, 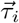, at position x_i_ in the filament is used to determine the position of the next structural unit in the chain as

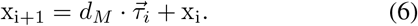

With *d*_*M*_ the distance between two consecutive structural units. Filament centerlines are modelled as helical curves aligned along the Z-axis so the tangent at a point is computed as

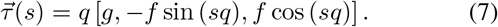

Where *s* is the length of the curve from its origin to the position of the point and *q* = (*f* ^2^ + *g*^2^)^−1^. The parameters of the helix curve are the radius *f* and the slope *f*/*g*. These parameters can be assigned from a given persistence length *p*, the physical parameter used to characterize polymer rigidity [39], using the following expressions

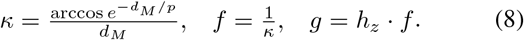

Where *κ* is the curve curvature in the XY-plane and *h*_*z*_ *∈* [0, 1] determines the curve elevation along the Z-axis as a fraction of the curvature, thus giving the curve also a torsion. After being placed on the filament curve, each structural unit can be additionally rotated around the tangent of the curve by an angle of

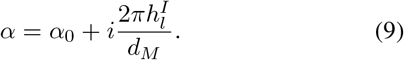

With *α*_0_ an arbitrary starting angle to allow randomization, *I* the position of the structural unit on the filament, and 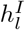 the length on the curve required to complete a turn. Consequently, we have two helices, the curve, or centerline, where the structural units are placed to give curvature and torsion, and the helix formed by the angle between two consecutive structural units, which model a possible helical inner structure of the filament as for actin. After defining the curve used to model the filament center-line from a given persistence length, the different structural units are inserted on the centerline. Below we present the models for the structural units for the two types of filaments used so far:
  - **Microtubules (MT):** the structural unit is modelled as a set of spheres with a radius of 40Å . Structural units are composed of 13 smooth-edged 3D solid spheres placed in a circle of radius 125Å on the XY-plane every 2π/13 radians, thus forming a complete circle as described in [40]. The distance between two consecutive structural units is set to 60Å, which is 1.2 times the unit length in the Z-axis (sphere diameter). MTs do not have a helical inner structure so the turn angle is *α* = 0.
  - **Actin-like:** the structural unit is modelled as a pair of spheres with a radius of 25Å aligned with the X-axis and separated by 50Å . The distance between two consecutive units is set to *d*_*M*_ = 60Å, which is 1.2 times the sphere diameter. The turn angle is *α* = 2π/12.5 because 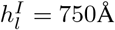 . This model approximates and averaged actin filament as described in [41].
3. *Macromolecules:* Cellular macromolecules can float in the cytoplasmic matrix or be bound to membranes. They can also be associated with filaments, but this case is not yet considered here. In addition, macromolecules can be randomly distributed or aggregated, creating clusters. Here we use the self-avoiding worm-like chain (SAWLC) model for polymers [42] to create these clusters. The SAWLC model also allows for random distribution of macromolecules if the polymer size is equal to that of a single macromolecule. Unlike the filamentous polymers described above, we assume here that the polymers have no rigidity and thus the clusters have no specific shape.

**Cytosolic:** in the SAWLC model each structural unit (or macromolecule), *M*_*i*_, is placed at a random point of the next set

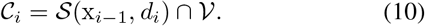

Where *d*_*i*_ is a variable distance from the previous macro-molecule *M*_*i*−*i*_ in the sequence, and 𝒮 a sphere centered at position, x_*i*−1_, and radius *d*_*i*_.

**Membrane-bound:** in the SAWLC model each macro-molecule, *M*_*i*_, is placed at a random point of the next in the next set

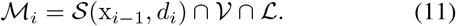

In comparison with cytosolic macromolecules, membrane-bound proteins must also be embedded in a membrane set (see Section II-A.1 Equation 5) ℒ.

**Heterogeneous clusters:** a SAWLC cluster, either cytosolic or membrane-bound, can contain *J* different types of macro-molecules, *M* ^*j*^, with *j ∈* [1, *J*] *⊂* ℕ.

### B. Stochastic Transformation Models

Each new inserted structural unit has a specific rotation, ℝ, with respect to the reference structural model and a specific translation, *T*, to its final position in *D*. Depending on the structure type, the rigid transformations *R* and *T* are generated according to different stochastic models (transformation generators *G*^*T*^), thus generating a representative variety of structures with different positions and rotations. Similarly, the geometric parameters for membranes and filaments are also generated using stochastic models (structural generators *G*^*S*^).

1. *Membranes:* Translations and rotations are generated in the same way for all membrane varieties. Translations are expressed with a coordinates vector generated according to the complete spatial randomness (CSR) model [43], [44]. Rotations are expressed as unit quaternions and are generated using the algorithm described in [45] to approximate a uniform random distribution. The membrane thickness *t* and layer thickness *σ*_*l*_ parameters are obtained from a random uniform distribution in a specified range. To ensure that all generated thicknesses are always within a valid physical range [46], [47], the distribution limits must lie within [2.5, 4.5] nm. **Sphere:** the radius *r* is generated from a random uniform distribution within the following range: the minimum corresponds approximately to the smallest vesicles in cells (typically 75Å) and the maximum with the diameter of 𝒱. **Ellipsoid:** the lengths of the semi-axis a = (*a*_*x*_, *a*_*y*_, *a*_*z*_) are taken from random uniform distributions within the same range as the spheres. In addition, the eccentricities are defined as

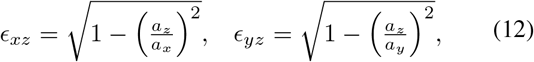

and must be smaller than a given value. **Toroid:** the two radii are obtained using a uniform random distributions similarly to sphere radius, then the greater value is assigned to the major radius *d* and the lower to the minor radius *e*.
2. *Filaments:* Unlike membranes, which are independent structures, filaments consist of a sequence of several structural units. Consequently, the rigid transformations for the structural unit, *M*_*i*_, depend on those applied to the previous unit, *M*_*i*−1_. Therefore a random generator for rigid transformations is only required for the first structural unit, *M*_0_, using the same distributions used for membranes.

For the filament geometry expressed as a helical curve, the only free parameters are the persistence length *p*, which controls the circular radius (and curvature) in the XY-plane, and the elevation in the Z-axis as a fraction of the circular radius *h*_*z*_, which controls the torsion. The values of *p* are derived from a random exponential unit distribution added to a minimum value of persistence length, this value sets a _1_maximum to the filaments’ flexibility. The values of *h*_*z*_ are obtained from a random uniform distribution between a given value (≤ 1) and 1.

In the case of actin-like filaments, it is possible to create a branched network, instead of generating isolated filaments. In a branched network, each new filament may start from any location in the VOI, like regular filaments thus starting a new network, but with a certain probability, *P*_*b*_ *∈* [0, 1], it starts from a random point of an already inserted filament, creating a branch within an already existing network. These parameters thus control the average number of networks to be generated for a given maximum occupancy and the branching density (number of branches per network length).

### C. Algorithms

This section contains the description of the different algorithms used to insert the structures into the VOI, *V*. First, the order chosen to process each structural type seen above is listed. Then, the algorithms for inserting each specific structure are described.

1. *Structure type selection:* Algorithm 1 details the procedure for processing the structural models. Only after all the structures of one type have been inserted, then we move to the next type. In each type, the order of each entity, e.g. membrane geometry (spherical, ellipsoidal or toroidal), is an input that can be controlled by the user. The criterion used to establish the order specified in Algorithm 1 is to place potentially larger and more rigid structures first. We assume that a cluster of small macromolecules following the flexible SAWLC model will be inserted more easily than a membrane or a filament in a highly fragmented volume.

#### Algorithm 1 Order of insertion for the different types of structures.

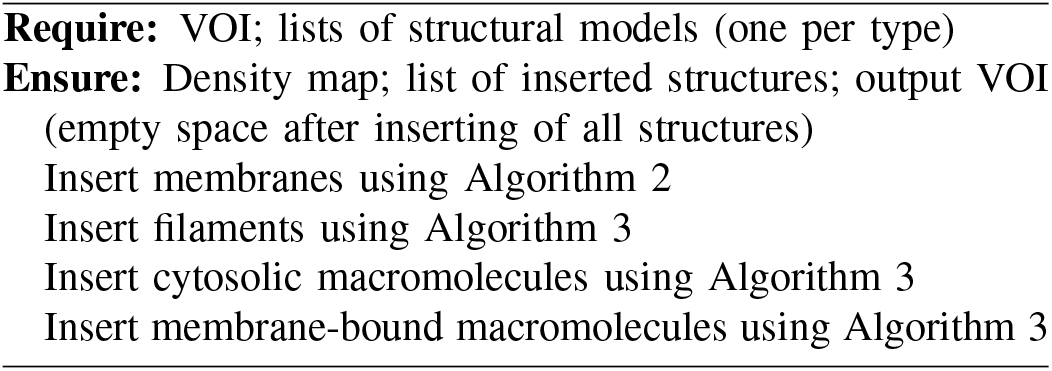
2. *Structure insertion:* The procedure for inserting membranes is described in Algorithm 2, for filaments and macro-molecules this procedure is identical and explained in Algorithm 3. Both algorithms are based on the trial-and-error principle: one tries to insert a new instance of a structure according to some geometrical constraints defined by the structural generator *G*^*S*^ into a space defined by the transformations generator *G*^*T*^, if this is not possible due to overlap with already inserted structures or because a fraction of the structure is outside of the VOI, then the algorithms try again by generating new geometric and transformation parameters. An additional stopping criterion is added to ensure the end of the process after a certain number of failed tries. Information about each structure is entered in the form of an input list *Q* including: the structural unit *M*, the target occupancy *O* and the statistical models for generating the structural parameters *G*^*S*^ and the rigid transformations *G*^*T*^ . Occupancy is the percentage of a volume occupied by one or more structures. In Algorithm 2, for each membrane model *i* in the list *Q*, an instance *M* with parameters generated by *G*^*S*^ is inserted into the updated VOI, 𝒱 _*o*_, according to the rigid body transformations *R* and *T* generated by the transformation generator *G*_*T*_ . The insertion loop for a membrane model *i* ends when the target occupancy *O*_*i*_ is reached, and then the algorithm proceeds to the next membrane model in the list *i*+1 until all models have been processed. The input 𝒱 is updated for each inserted membrane to exclude the already occupied volume. Therefore, 𝒱 _*o*_ will define the final empty space and corresponds to the VOI input for the next algorithm, which will process the next structure type.

#### Algorithm 2 Insertion of membranes in the VOI

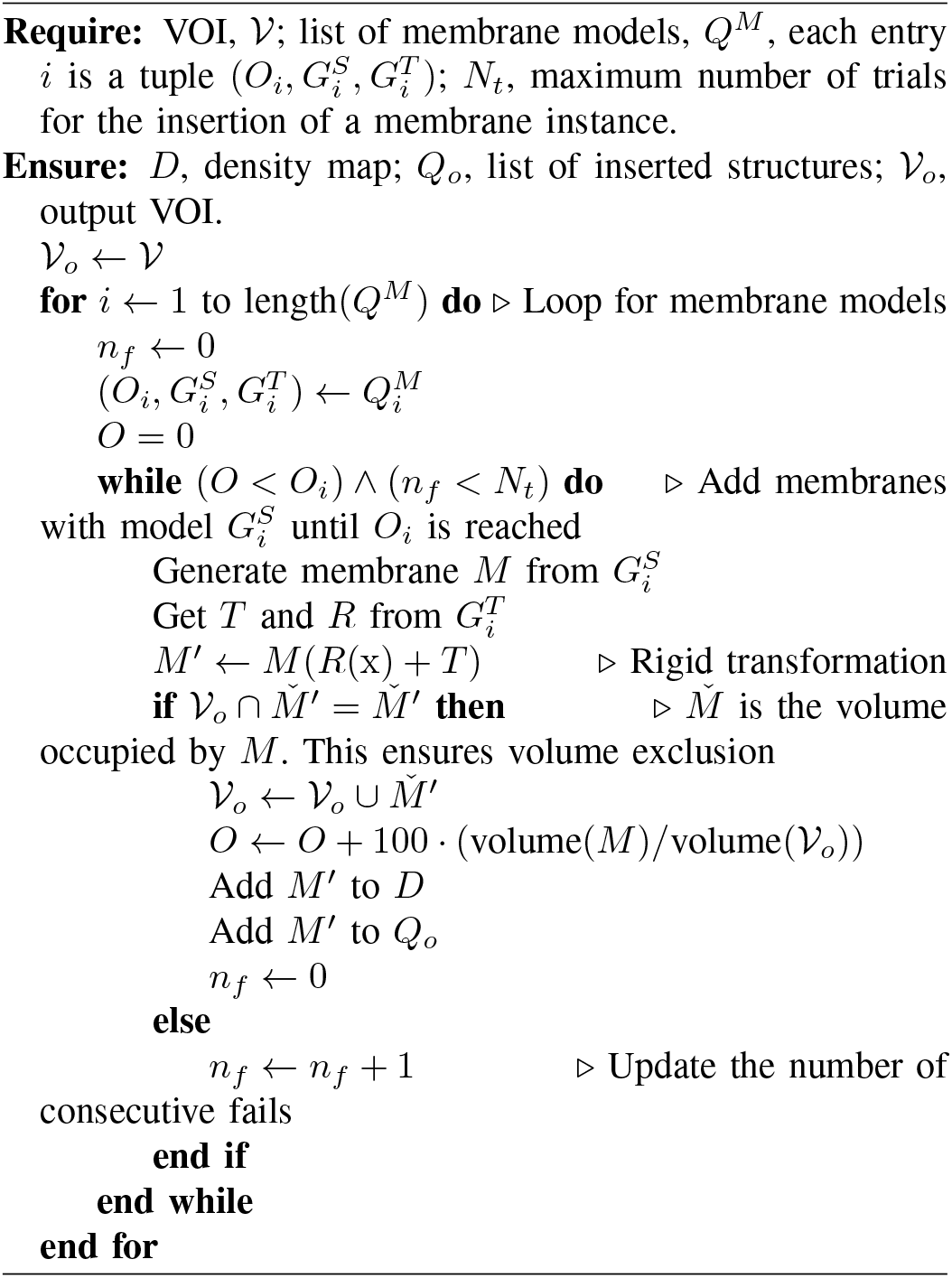 Regardless of overlapping, membrane instances are independent of each other. However, for filaments and macromolecules the insertion of the next structural unit depends on the previous one, except for the first one. Algorithm 3 is an extension of Algorithm 2 to handle this type of situation. It thus includes three nested loops. The first one iterates over all the introduced structural models. The second one attempts to reach the requested occupancy by inserting structural instances of a filament or cluster. The third one grows the filament or cluster by adding structural units. Randomization in Algorithms 2 and 3 is introduced through the stochastic structural and transformation models *G*^*S*^ and *G*^*T*^ . The addition of the counter, *n*_*f*_, in Algorithm 2 and 3, forces a restart of the structure insertion loop after *N*_*t*_ failures. As shown in Section III-C that helps to achieve high concentration of structures and avoids excessively long running times.

#### Algorithm 3 Insertion of filaments or macromolecules in the VOI

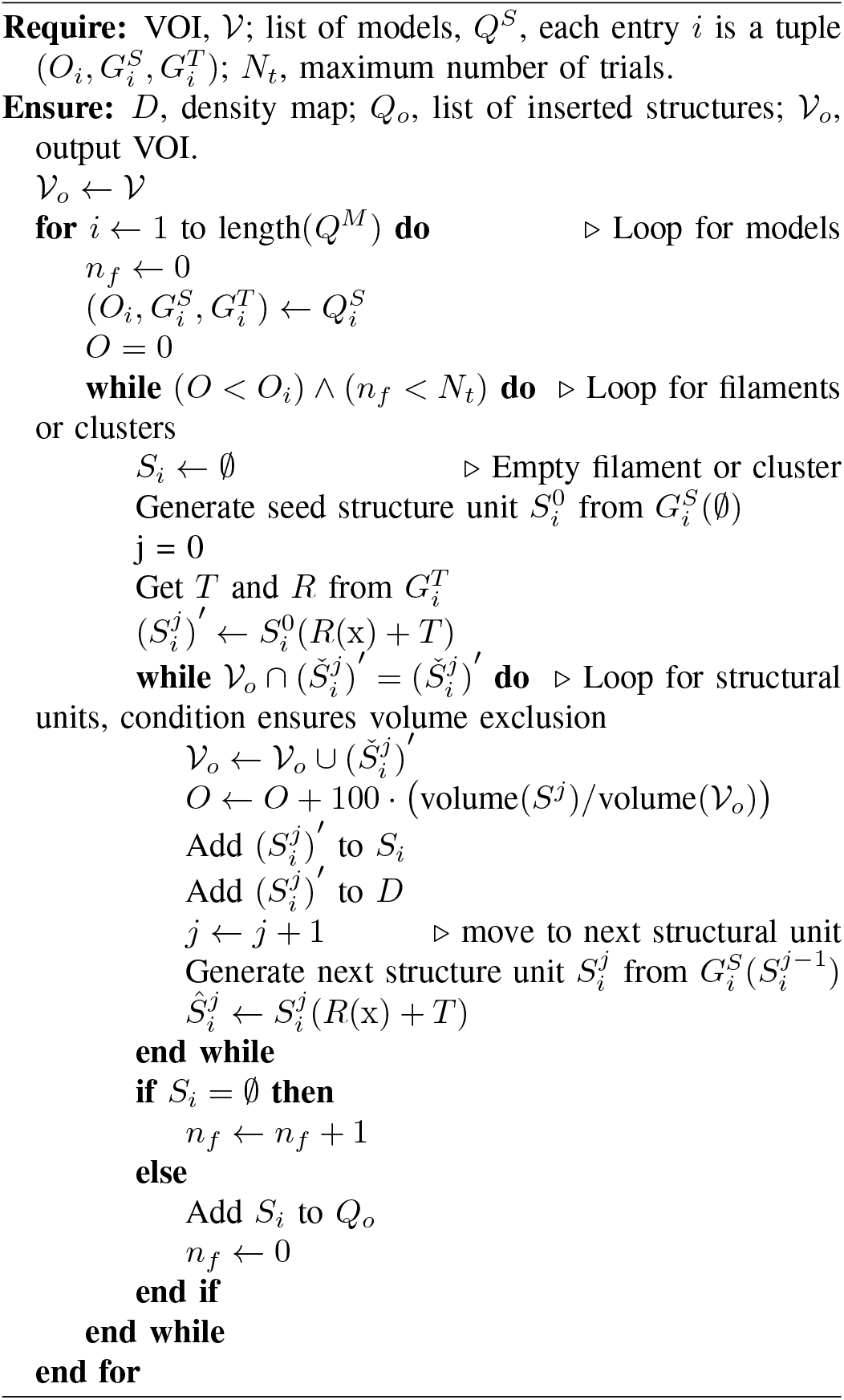

### D. Implementation details

Although this paper is dedicated to the simulation of cellular structures rather than the reproduction of the 3D image acquisition process in TEM, the implementation presented here is capable of reconstructing 3D tomograms containing some of the basic distortions inherent to cryo-ET such as noise, micrograph misalignment and missing wedge. The micrographs are generated as 2D projections at specified tilt angles using IMOD [48]. The noise is modeled by a Gaussian distribution, which is added to the micrographs, in order to approximate the tomogram contrast for a certain signal-to-noise ratio (SNR). Since the ground truth is known, noise level is adjusted to achieve certain SNR. Target SNRs are randomly generated from a uniform distribution within a specified range. The misalignment of the micrographs is modeled by applying random offsets on the X and Y axes of the micrographs in a sinusoidal model to penalize high tilt angles. Finally, the 3D tomogram is reconstructed from the micrographs using IMOD.

There are more accurate tools than IMOD for reconstructing cryo-tomograms from synthetic data [27], [29]. They have more complex models for noise, which also take into account solvent contribution. Since simulating the TEM process is still an active research field, here we generate enough output information (shape, position and rotation for each structural unit) to feed a TEM simulator. For example, very recently a software based on generative neuronal networks has been published to reconstruct tomograms from the synthetic micrographs [36] having distortions with realistic appearance.

## III. Results

Here we present the code implementation and synthetic data generated to show the potential of the proposed approach. All data were generated using a workstation running Ubuntu 22.04 LTS with an Intel^R^ Core™ i7-11700 2.50GHz processor, 128GB RAM and a GeForce RTX™ 3090 Ti GPU.

### A. Code

The software implementation of the models and algorithms described here is in a Python 3 package called ‘PolNet’ open-source available in the public repository *https://github.com/anmartinezs/polnet.git*. The scripts and settings for generating the next results are included in this repository.

### B. Cellular cryo-ET

The goal of this experiment is to simulate an exemplary tomograms with all currently available types of structures: membranes (spherical, ellipsoidal and toroidal), cytoskeleton (actin-like networks and microtubules), cytosolic and membrane-bound macromolecules. The size of the generated tomograms is 1000x1000x250 voxels, with a voxel size of 10Å, which is the typical configuration used today for tomogram visualization, molecular localization, and segmentation in cryo-ET.

Fig. 1.a shows an example of a generated clean density, free of noise and missing wedge. A ground truth semantic segmentation is also generated, the equivalent information is also stored in VTK format (see Fig. 1.b), which is more suitable for 3D visualization. In addition, the skeletons of the structures are also stored in VTK format, with filaments represented by their centerlines and clusters of macromolecules represented by a sequence of points connected by lines, with single points consequently corresponding to single macromolecules.

**Fig. 1.**
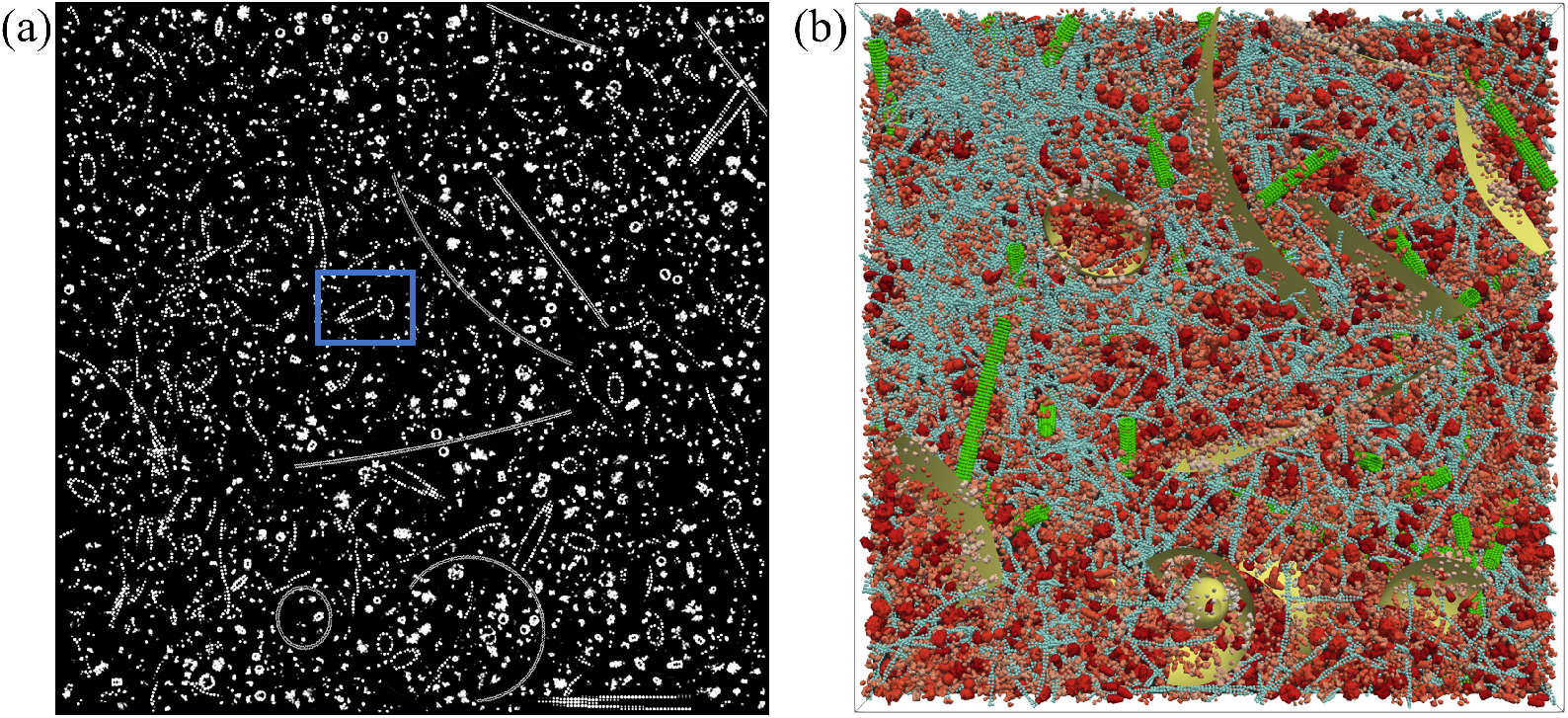
Simulation of the cellular context. (a) A 2D slice in the XY-plane of the generated density map, the blue box shows an example of how the growth of a microtubule was stopped to avoid overlap with another structure. (b) Polygonal ground truth data, membranes in yellow, actin-like network in blue, microtubules in green, macromolecules (included membrane-bound) in different shades of red.

Fig. 2.a shows a 3D reconstructed tomogram from 2D projected micrographs (*±*60°) that were randomly misaligned using a sinusoidal random model to penalize high tilt micrographs, with a mean value of 1 and a maximum misalignment of 1.5 pixels. As a result, the reconstructed tomogram suffers from distortions due to angular sampling and missing wedge. Gaussian noise was added to the micrographs to simulate the low contrast of the cryo-ET data by setting a target SNR randomly in the range [1, 2]. Fig. 2.b shows the missing wedge effect by comparing the reconstructed tomogram with the ground truth. The missing wedge effect is particularly visible in the YZ-slice where the membrane disappears and the macro-molecules are elongated on the Z-axis. The reconstructed and ground truth tomograms can be used as input information to train/test machine learning algorithms for image segmentation.

**Fig. 2.**
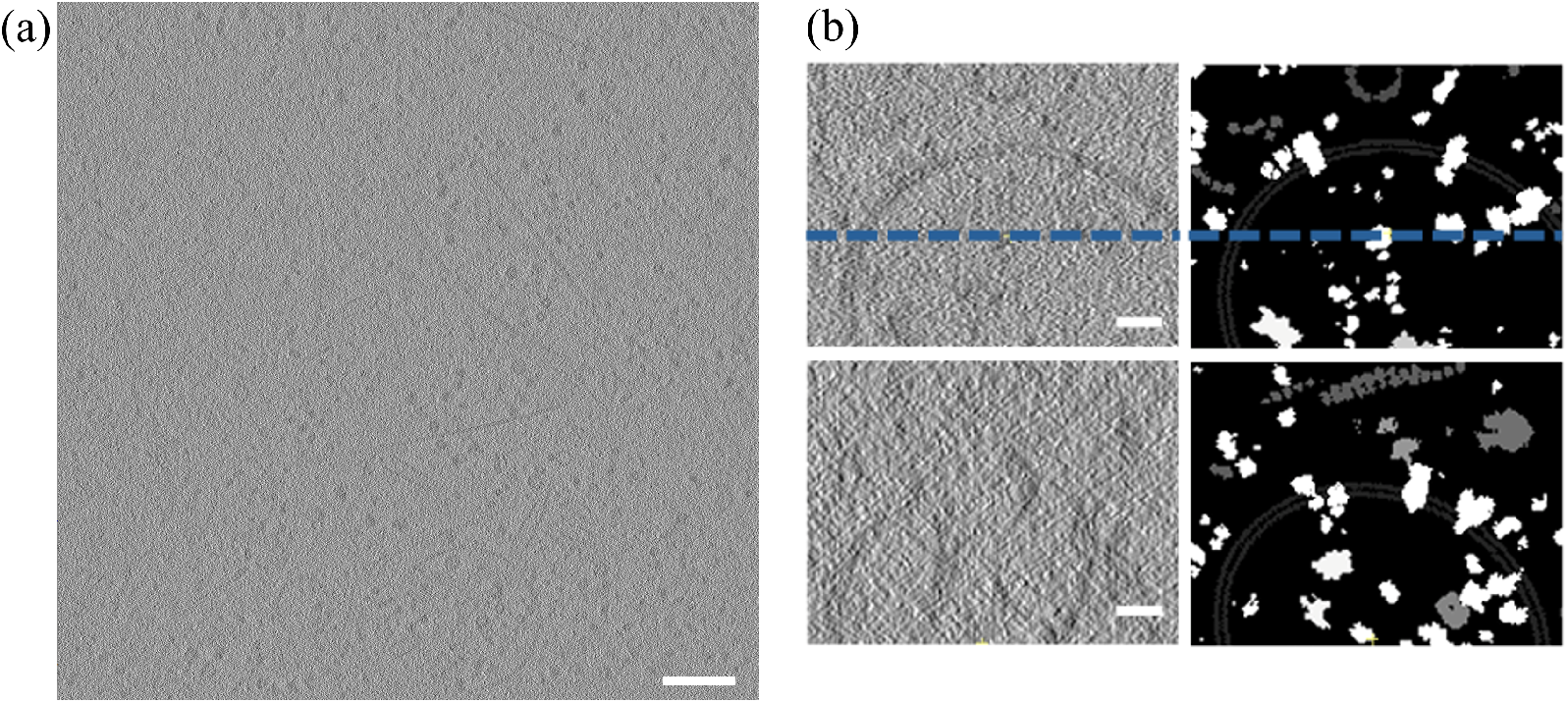
Reconstruction of tomograms. (a) A 2D slice in the XY-plane of a reconstructed tomogram corresponding to the density shown in Fig. 1. Scale bar 200 nm. (b) Zoom of the XY-plane showing membrane-bound proteins in the upper left, the label map is in the upper right, the dashed lines show the intersection with the YZ-plane slices shown in the bottom row. In the reconstructed YZ-slice (bottom-left) the membrane disappears at the apex and the macromolecules are elongated in the Z-axis relative to the ground truth (bottom-right) due to the missing wedge distortion introduced during reconstruction. The bilayer profile of the membrane can be appreciated in the magnified images. Scale bars 20 nm.

This example shows how Algorithm 1 can merge the different types of structures to generate a complex environment, which resembles the low order structures present in the cellular context.

### C. Simulation of the crowded environments

The cell cytoplasm and membranes usually have a high concentration of structures. Since structure insertion is based on a trial-and-error algorithm, the execution time increases with the concentration or occupancy. To evaluate structural insertion algorithms, here we run experiments to evaluate the insertion of cytosolic macromolecules, membrane-bound proteins and filaments separately. The membrane insertion algorithm is not evaluated because the total membrane occupancy generally does not exceed 1%. In these experiments, all tomograms had a size of 500x500x200 voxels, with a voxel size 10Å .

Regardless of the type of structure, the input occupancy is a target value. A trial-and-error algorithm cannot guarantee convergence within a certain time, which is especially difficult when the occupancy is very high. For macromolecules, Algorithm 3 terminates after failing *N*_*t*_ consecutive times to initialize a new structure. Increasing *N*_*t*_ therefore allows to obtain tomograms with higher densities at the cost of higher running times due to a greater number of structure insertion failures (see Fig. 3.a-c). An *N*_*t*_ = 100 achieves a target occupancy of 25% for cytosolic macromolecules (see Fig. 3.d-f), which already corresponds to a very crowded tomogram. Increasing *N*_*t*_ = 1000 allows to reach nearly 30% at the cost of approximately doubling the time.

**Fig. 3.**
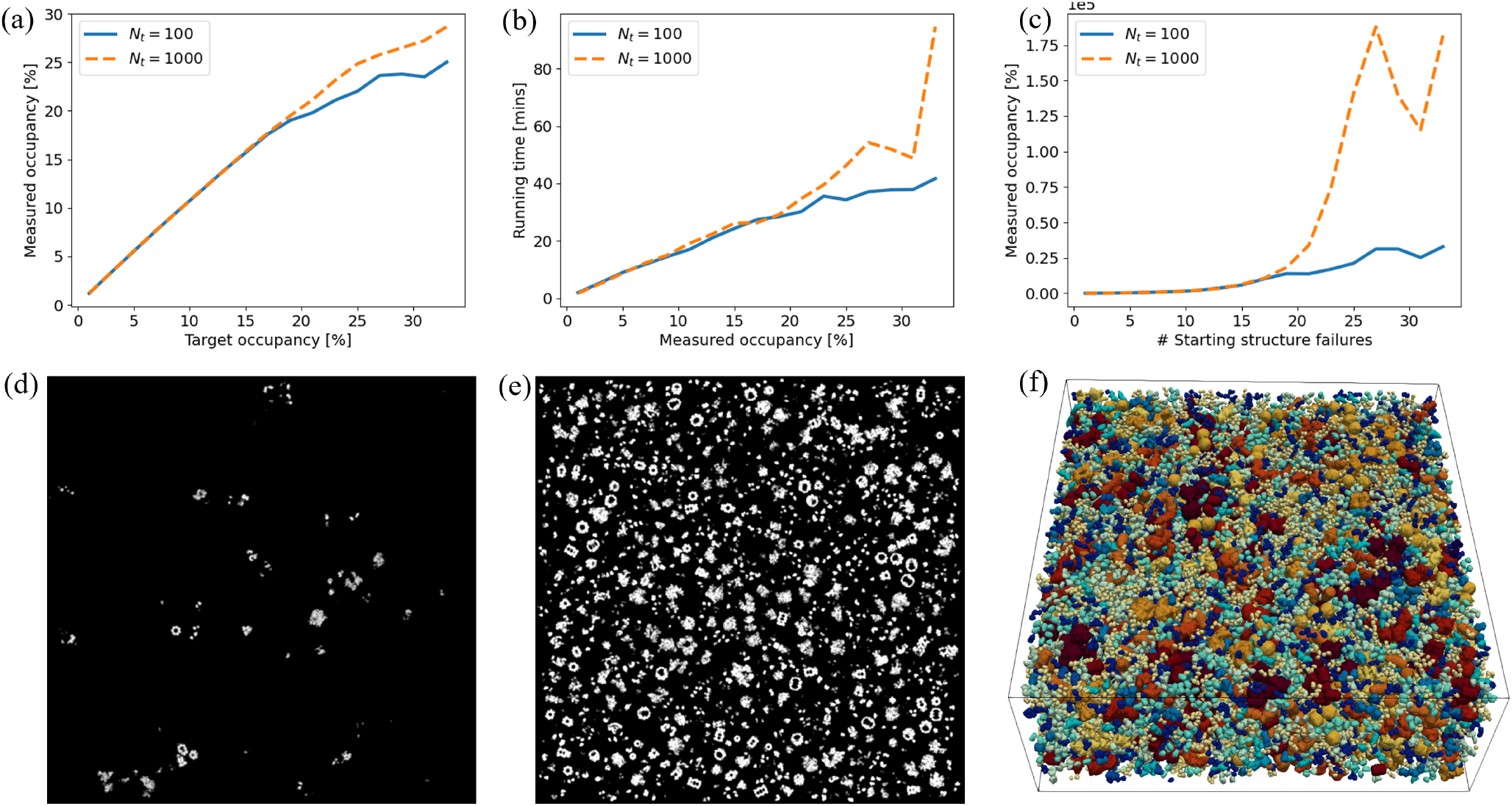
Simulation of cytosolic macromolecules. (a-c) Macromolecule insertion performance for different target occupancies where two ***N***_***t***_ values are evaluated. (a) Occupancy measured as the fraction of voxels belonging to macromolecules relative to the VOI volume. (b) Execution times. (c) Number of unsuccessful attempts to start a new macromolecule cluster. (d) 2D slice of a tomogram with **1%** occupancy. (e) 2D slice of a tomogram with a **25%** occupancy. (f) 3D view of the tomogram with a **25%** occupancy.

In contrast to cytosolic macromolecules, for which occupancy is calculated as the volume fraction (in percent) of the occupied VOI, the target occupancy for membrane-bound macromolecules represents the maximum area fraction (in percent) of the membrane surface occupied, as membrane-bound macromolecules can only be inserted on the membrane surface. Therefore, the input occupancy does not match the measured volume occupancy (see Fig. 4.a), although their dependence is approximately linear. Nanodomains, or clusters, of membrane-bound proteins are evident at low occupancies (Fig. 4.b). It is also possible to generate membranes with a high density of membrane-associated proteins, as in synaptic vesicles, viruses or endoplasmic reticulum (see Fig. 4.c).

**Fig. 4.**
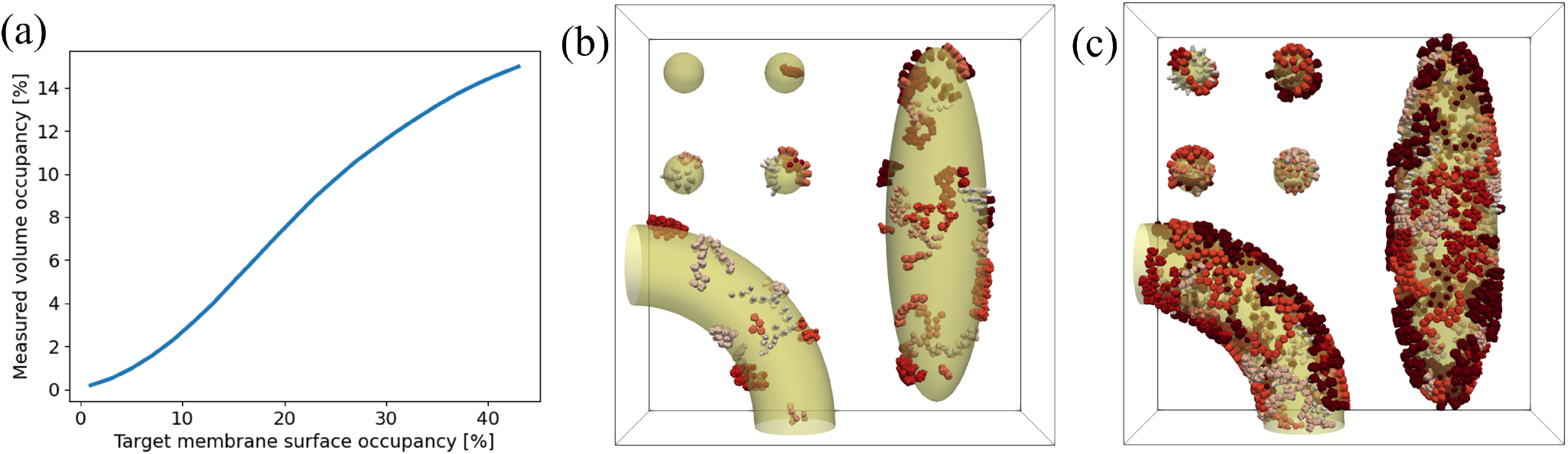
Simulation of membrane-bound proteins. (a) Measured output volume occupancy compared to the input target occupancy on membrane surfaces. (b) 3D visualization of a tomogram with a **7%** target occupancy for membrane-bound proteins. (c) Target occupancy of **39%**. (b-c) Membrane surfaces in transparent yellow, membrane proteins in different levels of red.

As for cytosolic macromolecules, the occupancy is computed from the filament volume. Occupancy values greater than 10% are not necessary because, in general, they do not correspond to realistic conditions in the cell. Fig. 5.b shows an occupancy of 7% which already results in a very dense actin-like network. An important parameter to control the shape of the filament networks is the branching probability for the filament *P*_*b*_ (see Fig. 5.a). The contrast between Fig. 5.b and 5.c shows that a high *P*_*b*_ value generates a densely concentrated filament network in specific regions, while conversely, a low *P*_*b*_ value generates a more evenly distributed network along the VOI.

**Fig. 5.**
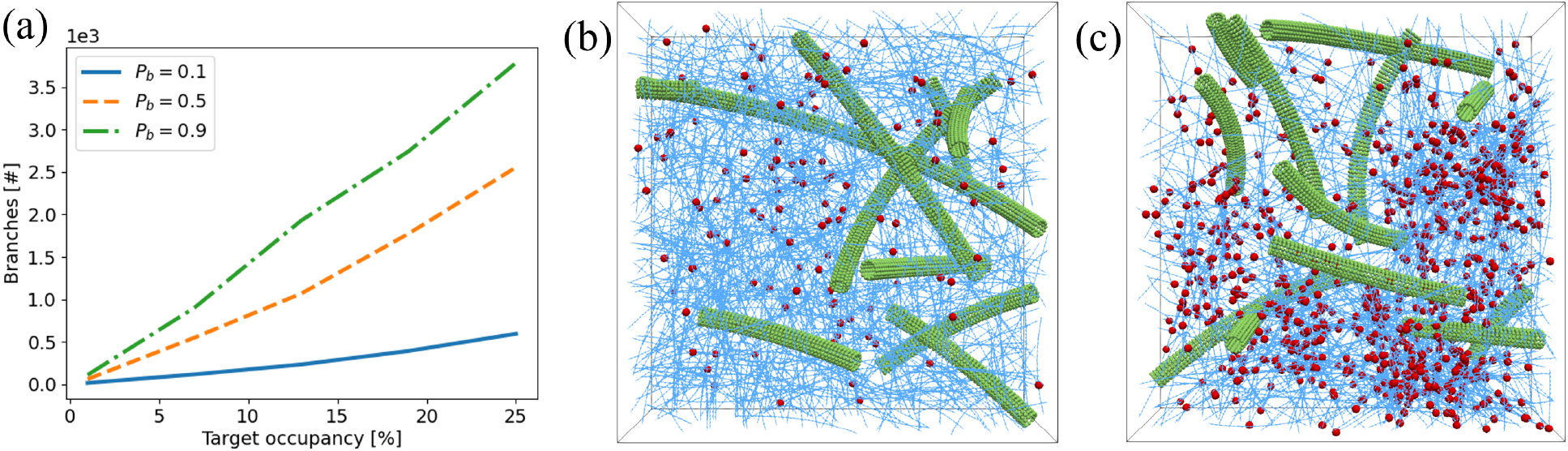
Simulation of filament-like structures. Target occupancy of **10%** for microtubules and **90%** for the actin-like filament network. (a) Number of actin-like filament branches for different occupancies and ***P***_***b***_. (b) 3D visualization of a tomogram with **7%** occupancy and ***P***_***b***_ **= 0**.**1**. (c) **7%** occupancy and ***P***_***b***_ **= 0**.**9**. In (b-c) only the centerlines are shown for actin-like filaments to allow a better visualization of the generated crowded network. Actin-like filaments in blue, actin-like filament branches as red spheres and microtubules in green.

### D. Training a deep learning model

To assess the usability of our synthetic data for training deep learning algorithms, we use a specific implementation of the U-Net architecture known as nnU-Net [49]. This framework is particularly known for its good performance without the need of manual hyper parameter tuning in the field of semantic segmentation for biomedical images. We trained nnU-Net to segment cryo-electron tomograms into the classes of membranes, microtubules, actin-like filaments, ribosomes, membrane-bound proteins and background (i.e., non of the other classes). To this end, we utilized the dataset presented in Subsec. III-B, and used ten reconstructed tomograms (see an example in Fig. 2.a) for training. For the model design, we chose nnU-Net 3D full resolution option, and the option fold all for training. We use the v2 of nnUNet software. As depicted in Fig. 6, the resulting prediction from our network validates our approach. Despite being trained solely on synthetic data, the network was capable of coarsely, but effectively, segmenting the main features of several *in situ* tomograms from different samples.

**Fig. 6.**
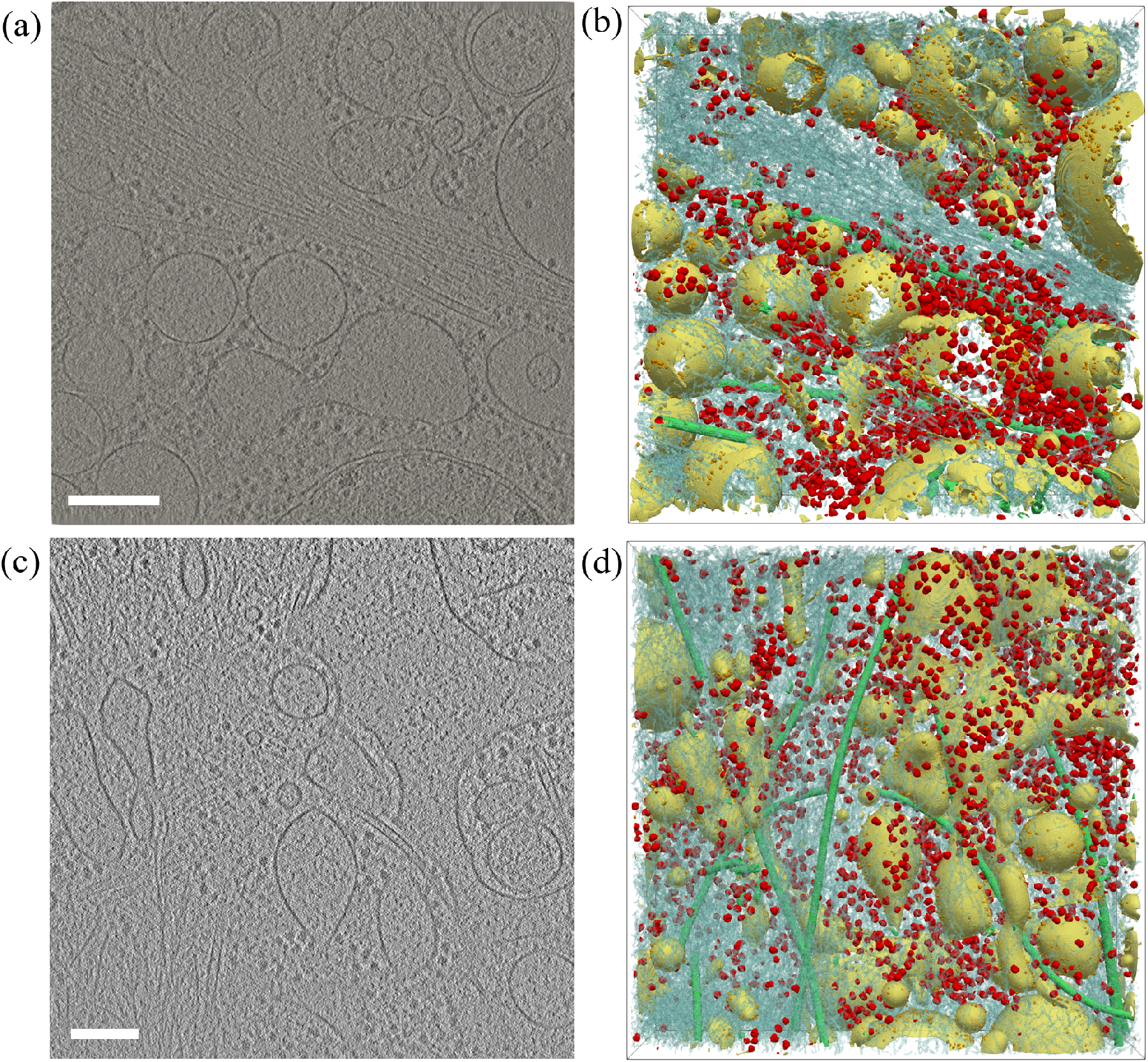
Training a deep learning model. (a) A 2D slice in the XY-plane of an *in situ* tomogram taken from EMD-10439. (b) Segmentation performed by nnU-Net trained solely from the synthetic data, membranes are in yellow, microtubules in green, actin-like filaments in transparent blue and ribosomes in red. (c-d) Similar to (a-b) but the tomogram was provided by [50] authors. It is remarkable the performance of our model to recover membrane areas faded out by missing wedge when compared with segmentation provided in the Fig. 1 of [50]. Scale bars 200 nm.

### E. Evaluation of the deep learning model

We have also processed tomograms providing as reference a segmentation to quantitatively validate the model generated in Subsection III-D. Specifically, we have taken publicly available tomogram EMD-11992, which is accompanied with segmentation for microtubules, actin filaments and ribosomes [51], and EMD-13671, with segmentation provided by authors [52] for intermediate filaments, actin filaments and membranes.

The model output is a semantic segmentation of the tomograms, therefore we should use Dice. However, as described in [53], the accurate segmentation evaluation for tubular structures such as filaments requires novel similarity metrics to preserve topological features meanwhile arbitrary features like thickness are neglected. Authors in [53] proposes clDice (centerline Dice) for tube-like structures in 2D images, here we extend this metric to 3D for processing filament like structures. Analogously, we define and compute sDice (surface Dice, described also in [54]) for membranes and pDice (point Dice) for macromolecules. Consequently, clDice variant is computed for microtubules, actin or any other filamentous structure, pDice for ribosomes because these are macromolecules, and sDice for membranes. These metrics are calculated from the previous computation of the topology precision (TP), the fraction of the predicted segmentation skeleton within the ground truth, and the topology sensitivity (TS), the fraction of the ground truth skeleton within the predicted segmentation. In this paper, all Dice values shown are computed from pDice, clDice or sDice depending on the processed structures as described above.

Table I contains the quantitative evaluation results obtained for the structures with reference segmentation in EMD-13671. For membranes, the segmentation agrees very well the reference. However, it is important to consider in this case the reference segmentation has been provided by a DL model, MemBrain-v2 [54], specifically designed for segmenting only membranes and trained with a large and diverse experimental dataset. More importantly, the high sensitivity value shows that our model segments almost everything predicted by MemBrain-v2, but as shown in Figure 7 our model seems to partially recover regions missed by missing wedge. Regarding the results for actin filaments, Dice scores are very low, but specificity values are very high. Additionally, Figure 8 shows that besides actin filaments, intermediate filaments are also segmented by our model. To further investigate the capacity of our model to become a general detector for multiple types of filament networks, we processed dataset EMD-13671 with two models: one trained with all features as described in Section III-D (left values in Table I), and a second without the model for actin-like filament networks but including a high resolution model (PDB-5MVY) of an straight actin filament fragment (right values in Table I). The second model does not improve significantly the precision but presents a worse sensitivity. Moreover, it mostly fails to recover intermediate filaments (see Figure 8.a-c). The comparison of these models reinforces the necessity of the filament network model presented here to train a general model for processing filaments with different inner structures.

**TABLE I.** Segmentation evaluation for EMD-13671.

|  | Dice | TP | TS |
| --- | --- | --- | --- |
| **Actin** | 0.48 / 0.43 | 0.34 / 0.36 | 0.75 / 0.54 |
| **Inter. Filament** | 0.05 / 0.02 | 0.02 / 0.01 | 0.68 / 0.25 |
| **Membrane** | 0.71 | 0.60 | 0.87 |

**Fig. 7.**
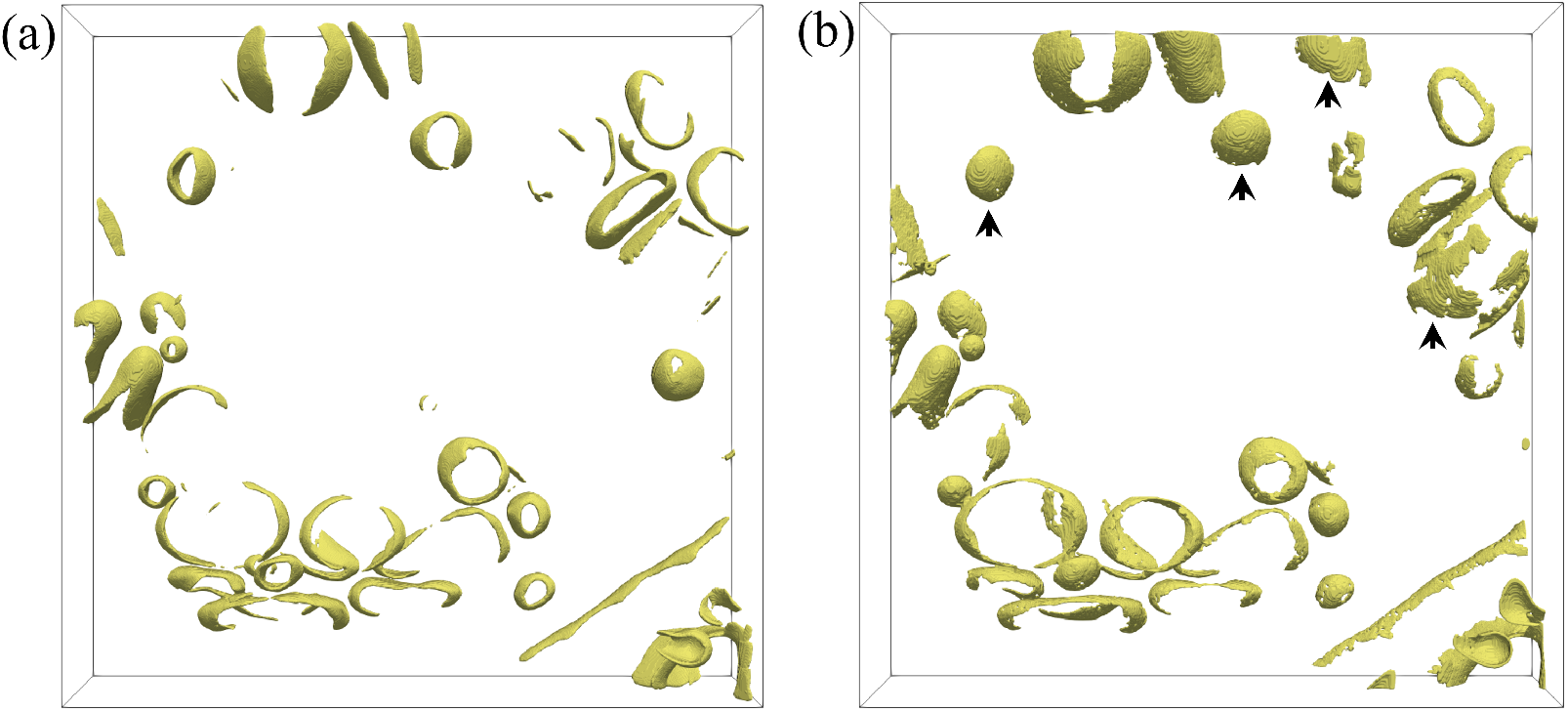
Membrane segmentation for EMD-13638. (a) Reference segmentation by MemBrain-v2 [54]. (b) Segmentation of our model, arrows depict top-view membranes partially recovered despite missing wedge.

**Fig. 8.**
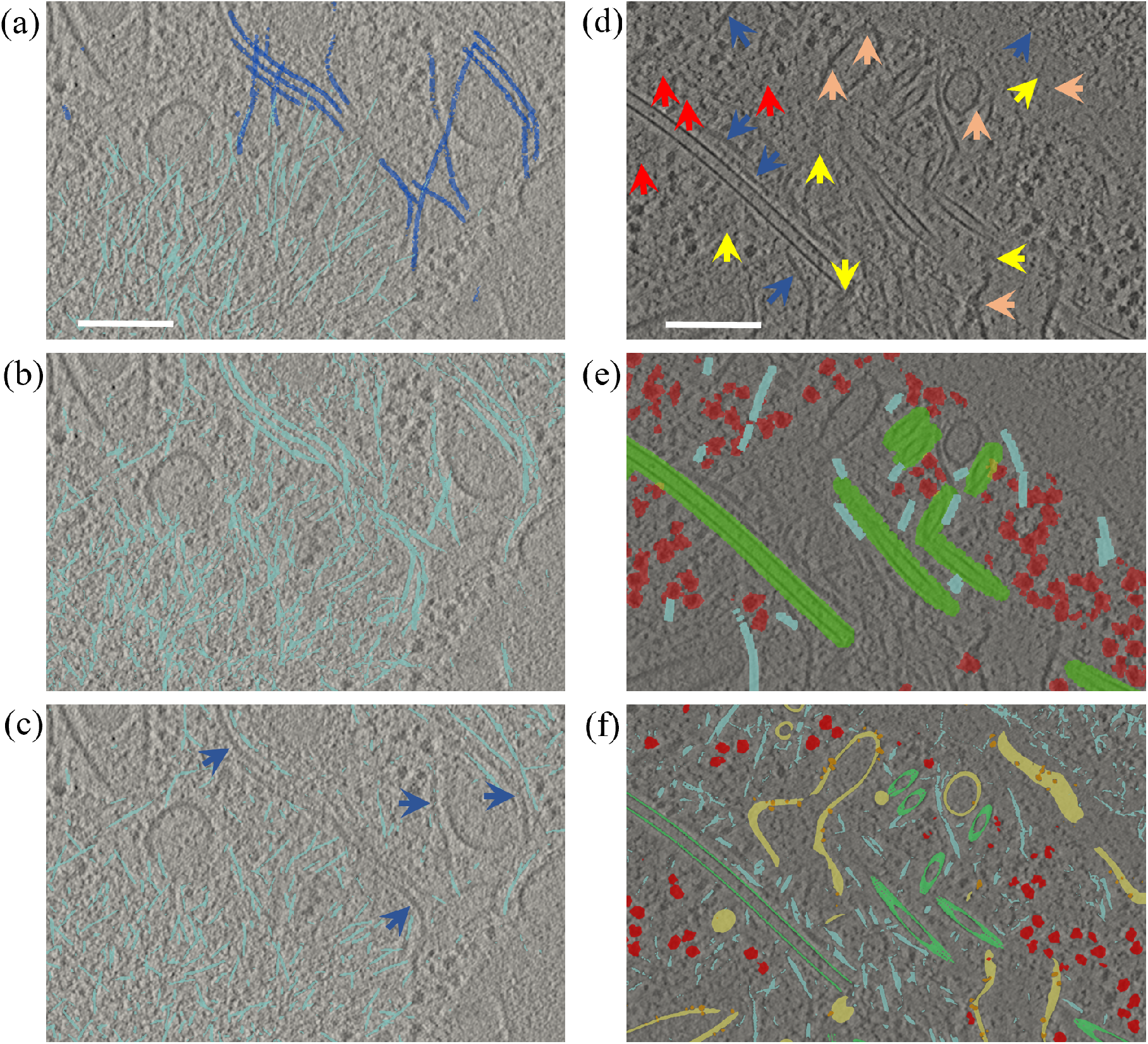
Detailed segmentation evaluation. (a-b) Filamentous networks in a 2D slice of EMD-13671. (a) Manually supervised reference segmentation provided by [52], light blue are actin-like filaments and dark blue intermediate filaments. (b) Segmentation by our DL model trained with synthetic data including actin-like networks. (c) The synthetic training set includes instances of a rigid model for actin (PDB-5MVY), but not the actin-like networks presented here. Dark blue arrows point to intermediate filaments missed, especially in areas of high curvature. (d-f) EMD-11992. (d) Blue arrows point to filamentous structures not analyzed in (e), yellow arrows show membrane top-views where the membrane bilayer vanishes (see (f)), red arrows show potential false positives for ribosomes in (e), and orange arrows point to some membrane proteins including top-views (see (f)). (e) Reference segmentation provided in EMD-11992, red ribosomes, light blue actin filaments and microtubules in green. (f) Segmentation obtained with our model, membranes in yellow and membrane bound proteins in orange. Scale bars 200 nm.

Table II contains the quantitative evaluation results obtained for the structures with reference segmentation in EMD-11992. The ribosome Dice score is around 0.5 with precision above 0.8. This is very interesting because the reference segmentation for ribosomes contains potential false positives depicted in Figure 8.a. Microtubules are almost perfectly segmented. Regarding actin, the situation is similar to EMD-13671, the sensitivity is high but the precision is low because the model segments many other filament-like structures besides actin filaments. Although EMD-11992 does not include a proper reference segmentation for membranes and membrane-bound proteins, Figure 8.d-f shows that our model performs well and is able to recover some top-view membranes and membrane-bound proteins, which are normally elusive because the lipid bilayer vanished due to the missing wedge.

**TABLE II.** Segmentation evaluation for EMD-11992.

|  | Dice | TP | TS |
| --- | --- | --- | --- |
| **Microtubule** | 0.95 | 0.97 | 0.93 |
| **Actin** | 0.19 | 0.10 | 0.89 |
| **Ribosome** | 0.51 | 0.82 | 0.37 |

## IV. Discussion

Although higher order structures such as organelles (mitochondrion, centriole, …) or the nucleolus would require specific and very sophisticated models, the simulation of low-order structures proposed here can already be used to study many interesting cellular processes such as membrane morphology [23], [24], filament interaction under pathological conditions [22], macromolecule clusters [17], macromolecules in the liquid-like state [19] and membrane-bound protein nanodomains [18], [55]. In addition, local feature simulation has been shown a path to overcome the current limitations of DL approaches for processing (segmentation, macromolecular localization, …) cryo-tomograms [32], [34]. However, validating tools analyzing experimental cryo-ET data is particularly challenging. For example, only few public datasets come with a segmentation, typically done by using several computer methods assisted by manual restoration. In practice, subtomogram averaging is used to verify the quality of particle positions: A resulting high-resolution structure indicates that many positions are correct. However, that does not guarantee a low number in false positives or false negatives. Therefore, quantitative results require a detailed interpretation.

Interestingly, a DL model trained with synthetic data was able to predict some membrane regions faded out by missing wedge. This result opens a new approach to solve the missing wedge problem in cryo-ET. If we are able to reproduce enough contextual information in the synthetic data, then a DL model may recover the missing information. For the case of membranes, we plan to improve membrane models with the intention of generating more complex membrane arrangements. Additionally, our approach also allows to detect top-views for membrane bound proteins, something that is elusive to other specific methods [55].

Another interesting finding is that our model is not specific for a single type of filaments like actin. This model does not replicate specific actin features, but instead, is able to generalize the generic shape of a filament, in addition to its local geometry changes (curvature) and connections with other filaments. As a consequence, we have empirically observed that it can be used to train DL algorithms for parsing generic filament-like structures. In addition to actin, it detects many other filamentous structures such as intermediate filaments, as well as new structures not yet processed. We plan to further investigate how to include higher resolution features in filament structural models to obtain more specific detectors.

Reproducing the high concentration of macromolecules in the cellular cytoplasm or bounded to membranes is a major challenge. So far, only trial-and-error algorithm has been used for that purpose. Considering that we insert macromolecules in clusters (nanodomains), we have extended such approach by adding counters to re-start the insertion and avoid to get stuck in tomogram regions without enough empty space. This solution has allowed to generate synthetic tomograms with a very high concentrations of macromolecules, similarly to the cellular context. Our plan is to investigate the parallelization of the algorithms to speed up simulations.

We demonstrate that our synthetic data approximate well many of the features present in experimental *in situ* tomograms. Specifically, the information contained in the synthetic data is sufficient to train a generalized DL model to segment main features present in real cellular tomograms. Results show that segmenting observable features such as membranes and microtubules, as well as, large macromolecules like ribosomes or some membrane bound complexes does not require subnanometric information. Consequently, no external software was required to simulate an accurate model for CTF modulation, although PolNet is prepared to interact with external TEM simulators. Nevertheless, as future work, we plan to include accurate in-house CTF modulation for tomograms to reproduce high resolution (sub-nanometric) features. We foresee such information will become critical to reproduce accurately sub-nanometric features, and these may be necessary to recognize small structures by DL.

## Acknowledgments

We thank Prof Dr Juergen Plitzko at the Max Planck Institute of Biochemistry for sharing computational resources needed for software development. We thank Tingying Peng from Helmholtz AI for mentoring L.L. We thank Jonathan Schneider for providing and segmenting tomogram in Fig 6.c.

